# Somatic point mutations are enriched in long non-coding RNAs with possible regulatory function in breast cancer

**DOI:** 10.1101/2021.07.19.453012

**Authors:** Narges Rezaie, Masroor Bayati, Maedeh Sadat Tahaei, Mehrab Hamidi, Sadegh Khorasani, Nigel H. Lovell, James Breen, Hamid R. Rabiee, Hamid Alinejad-Rokny

## Abstract

De novo somatic point mutations identified in breast cancer are predominantly non-coding and typically attributed to altered regulatory elements such as enhancers and promoters. However, while the non-coding RNAs (ncRNAs) form a large portion of the mammalian genome, their biological functions are mostly poorly characterized in cancers. In this study, using a newly developed tool, SomaGene, we reanalyze *de novo* somatic point mutations from the International Cancer Genome Consortium (ICGC) whole-genome sequencing data of 1,855 breast cancers. We identify 929 candidates of ncRNAs that are significantly and explicitly mutated in breast cancer samples. By integrating data from the ENCODE regulatory features and FANTOM5 expression atlas, we show that the candidate ncRNAs in breast cancer samples significantly enrich for active chromatin histone marks (1.9 times), CTCF binding sites (2.45 times), DNase accessibility (1.76 times), HMM predicted enhancers (2.26 times) and eQTL polymorphisms (1.77 times). Importantly, we show that the 929 ncRNAs contain a much higher level (3.64 times) of breast cancer-associated genome-wide association (GWAS) single nucleotide polymorphisms (SNPs) than genome-wide expectation. Such enrichment has not been seen with GWAS SNPs from other diseases. Using breast tissue related Hi-C data we then show that 82% of our candidate ncRNAs (1.9 times) significantly interact with the promoter of protein-coding genes, including previously known cancer-associated genes, suggesting the critical role for candidate ncRNA genes in activation of essential regulators of development and differentiation in breast cancer. We provide an extensive web-based resource (http://ncrna.ictic.sharif.edu), to communicate our results with the research community. Our list of breast cancer-specific ncRNA genes has the potential to provide a better understanding of the underlying genetic causes of breast cancer. Lastly, the tool developed in this study can be used in the analysis of somatic mutations in all cancers.

## Introduction

Despite the substantial early enhanced diagnostic techniques of breast cancer (BC) and the development of adjuvant therapy, radical and personalized surgery, breast cancer is still the most common cancer diagnosed among women. It is still the highest frequently leading cause of cancer-related mortality amongst females worldwide [1]. A deeper understanding of the underlying mechanisms of breast cancer genetics and pathogenesis can be used for detecting early-stage cancer cause to reduce morbidity and mortality as a result of breast cancer [2]. Over the last ten years, high-throughput sequencing has provided a comprehensive investigation on the underlying genetic mechanisms that initiate or drive cancer progression and revealed many cancer-associated mutations. International Cancer Genome Consortium (ICGC) [3] has provided an extensive catalog of somatic mutations for various cancer types. This information has enabled researchers to characterize numerous protein-coding genes that play an essential role in cancer progression [4–8].

Although most studies have mainly focused on protein-coding genes in investigating driver mutations in cancer, several lines of evidence imply that about 80% of the genome is biochemically functional. Additional to the lack of information on the genome’s non-coding regions, our current understanding of genetic information’s present functional consequences is highly primitive. It focuses instead on proteins found in only 2% of the genome. However, a large portion of non-coding regions operate as regulators of oncogenes [9] and non-coding RNAs (ncRNAs), which share traits with mRNAs such as sequence conservation, polyadenylation, and splicing [10, 11]. Additionally, around 93% of disease-associated genome-wide association single nucleotide polymorphisms (GWAS SNPs) are located within these regions [12, 13], which can significantly influence gene expression of coding and non-coding genes.

Long non-coding RNAs (lncRNAs), an influential class of non-coding transcripts with more than 200 nucleotides, are potential cancer progression indicators and are emerging as diagnostic biomarkers [10, 14]. For example, *GAS5* is one of the lncRNAs that is considerably down-regulated in breast cancer [15]. *HOTAIR* is another well-known lncRNA upregulated in breast cancer, contributing to aberrant histone H3K27 methylation and cancer metastasis [16, 17]. Furthermore, lncRNAs contribute to various regulatory activities in the cell, such as regulating gene expression via interaction with other chromatin regulatory proteins [18], function as active enhancers [19, 20], and regulate chromatin structure [21–23]. Despite various studies on the impact of non-coding somatic mutations occurring in ncRNAs, the role of such ncRNAs has remained underexplored in breast cancer.

In this study, we developed a new tool, SomaGene, to identify 929 ncRNAs that are significantly and specifically mutated in breast cancer. Using SomaGene, we show that candidate ncRNAs identified in our study are significantly enriched for regulatory features (e.g., breast-specific H3K27ac, H3K4me1, CTCF, DNase hypersensitive sites, and enhancer marks). Notably, we show that our breast cancer candidate ncRNAs have a much higher fraction of both GWAS SNPs and eQTL (expression quantitative trait loci) polymorphisms. Finally, our analyses on high throughput chromosome conformation capture (Hi-C) data from the Human Mammary Epithelial Cell (HMEC) indicate that many of our candidate ncRNAs significantly interact with at least one protein-coding gene which may suggest a potential enhancer role for these ncRNAs. An overview of the pipeline used in this study is shown in **Figure 1**.

**Figure 1.**
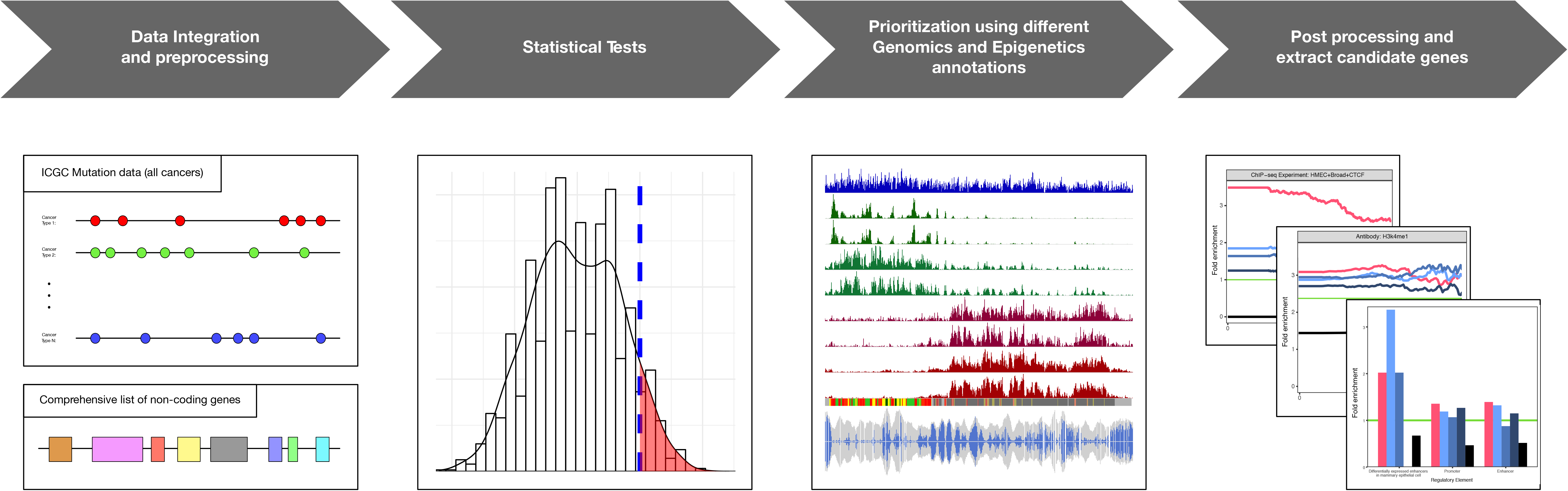
The flow diagram of the SomaGene pipeline used in this study. ICGC cancer samples are used to identified BC-associated non-coding RNAs. Only samples with single point mutations are considered in the analysis. We also used both the FANTOM5 robust gene list and Ensembl gene list to have a comprehensive list of non-coding genes. After counting the number of mutated samples in each ncRNA, we then used Fisher’s exact test described in the method section to identify mutational P-values for each ncRNA. To identify significantly mutated ncRNAs for each ncRNA, we calculated P-values for 1,000,000 random permutations of breast/non-breast labels to estimate the 99%C.I. threshold of P-values. We then investigated the overlapping of non-coding RNAs with breast tissue-related regulatory features (e.g., ENCODE predicted chromHMM, H3K27ac), BC-related GWAS SNPs, HMEC related eQTL polymorphisms, and HMEC related Hi-C interacting regions.

We also provide a list of non-coding genes with their mutational enrichment P-value and annotated genomic signals at http://www.ncrna.ictic.sharif.edu and https://www.ihealthe.unsw.edu.au/research. SomaGene as an open-source R package is also available at https://github.com/bcb-sut/SomaGene.

## Results

To have a comprehensive list of ncRNAs, we used a combined list of ncRNAs provided by the FANTOMCAT [24] and Ensembl [25] consortia (see method section). This includes ncRNAs from different types inclusive of pseudogene (22.9%), lncRNA intergenic (21.4%), long intergenic ncRNAs (5.6%), lncRNA divergent (13.4%), antisense (3.3%), lncRNA sense intronic (6%), miRNA (5%), misc RNA (3%), lncRNA antisense (4.8%). A full list of ncRNAs provided in **Supplementary figure S1a**. Somatic mutations from 19 cancer types were downloaded from ICGC [3], including 1,855 breast cancer samples containing 17,163,482 single point somatic mutations and 10,460 samples with other cancers containing 67,752,271 somatic point mutations inside the ncRNA regions.

### Background model to identify significant non-coding RNAs in breast cancer

To identify the ncRNAs that are significantly mutated in breast cancer samples, we counted the number of samples with somatic mutations in ncRNA from 1,855 breast cancer samples and 10,460 samples with other cancers. We then calculated a P-value for each ncRNA using Fisher’s exact test (*see method section*). To identify significantly mutated ncRNAs, we calculated P-values for 1,000,000 random permutations of breast/non-breast labels for each ncRNA and estimated the probability that an association emerges by chance (*see method section*). This provides a threshold for each ncRNA separately (**Figure 2**). As a result, we identified 929 ncRNAs (99% confidence interval – see method section) that significantly mutated in breast cancer samples (**Supplementary table S1**). Looking into our candidate ncRNAs revealed that 27.2% of them are lncRNAs intergenic, 18.1% pseudogene, 3.8% long intergenic non-coding RNAs (lincRNA), 20.8% lncRNA divergent, 2.5% antisense, and 6.6% lncRNA sense intronic (**Supplementary figure S1b**).

**Figure 2.**
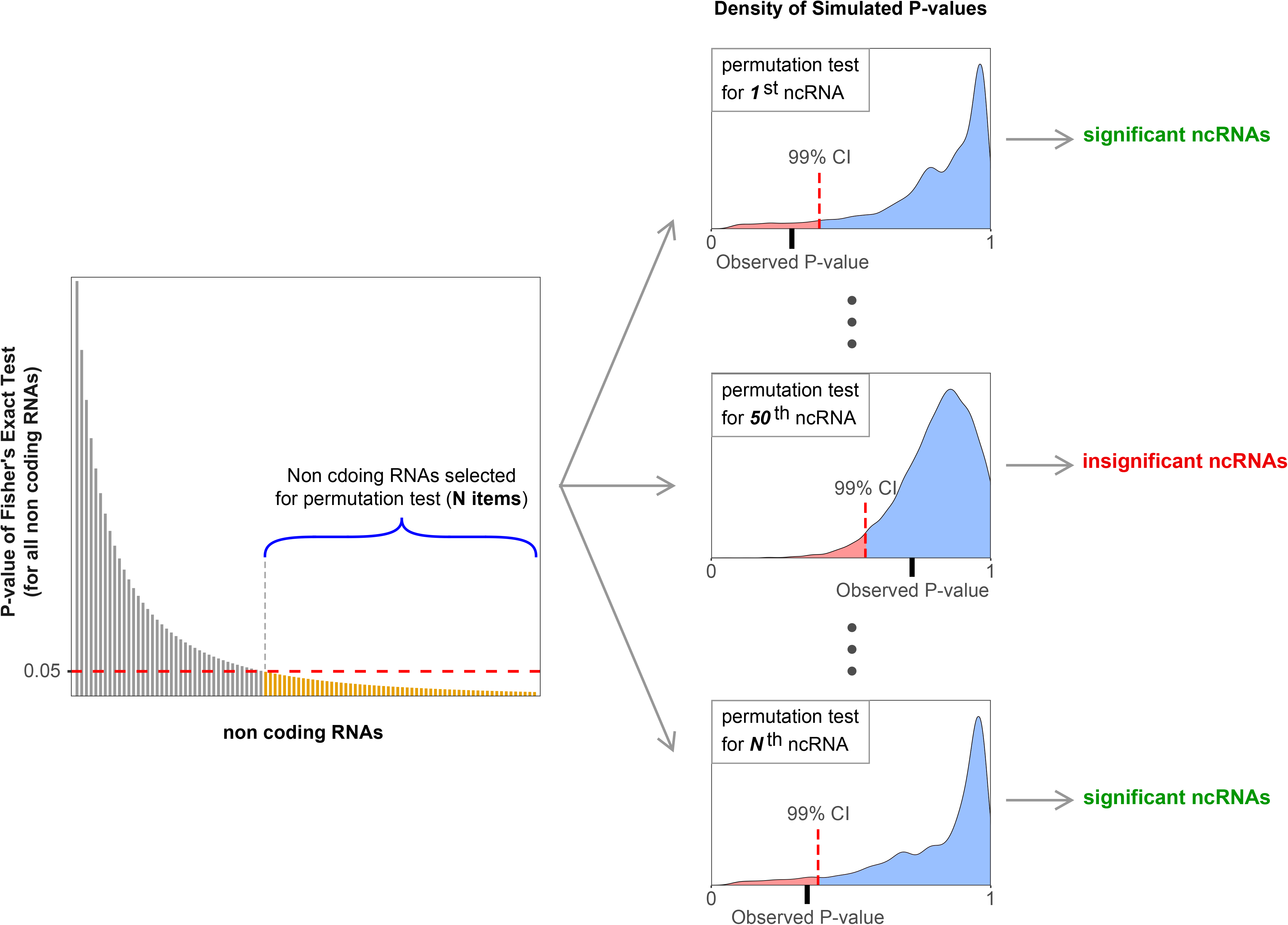
A schematic of permutation process to identify significantly mutated non-coding RNAs in breast cancer samples. To determine the significance level of mutations in ncRNAs, we calculated P-values for 1,000,000 random permutations of breast/non-breast labels to estimate the 99% C.I. threshold of P-values. The permutation accounts for statistical significance based upon our original data sampling and avoids Type II errors that may arise from multiple testing correction approaches such as Bonferroni. We performed the permutations for each ncRNA separately. In the figure, the red color indicates the 99% C.I. of observed P-values in the permutations.

### Enrichment of regulatory features in the BC-associated ncRNAs

To determine if the somatic mutations are enriched in ncRNAs with regulatory function, we first examined the enrichment of HMEC-related chromatin states provided by the ENCODE consortium within our significant ncRNAs. As **Figure 3a** shows, both ENCODE promoters and enhancers have been significantly enriched within our candidate ncRNA genes (3.23 and 2.26 times with P-values 4.12e-29 and 5.28e-32 enrichment for promoters and enhancers, respectively), suggesting breast cancer *de novo* somatic mutations are enriched in ncRNAs with enhancer and/or promoter like functions.

**Figure 3.**
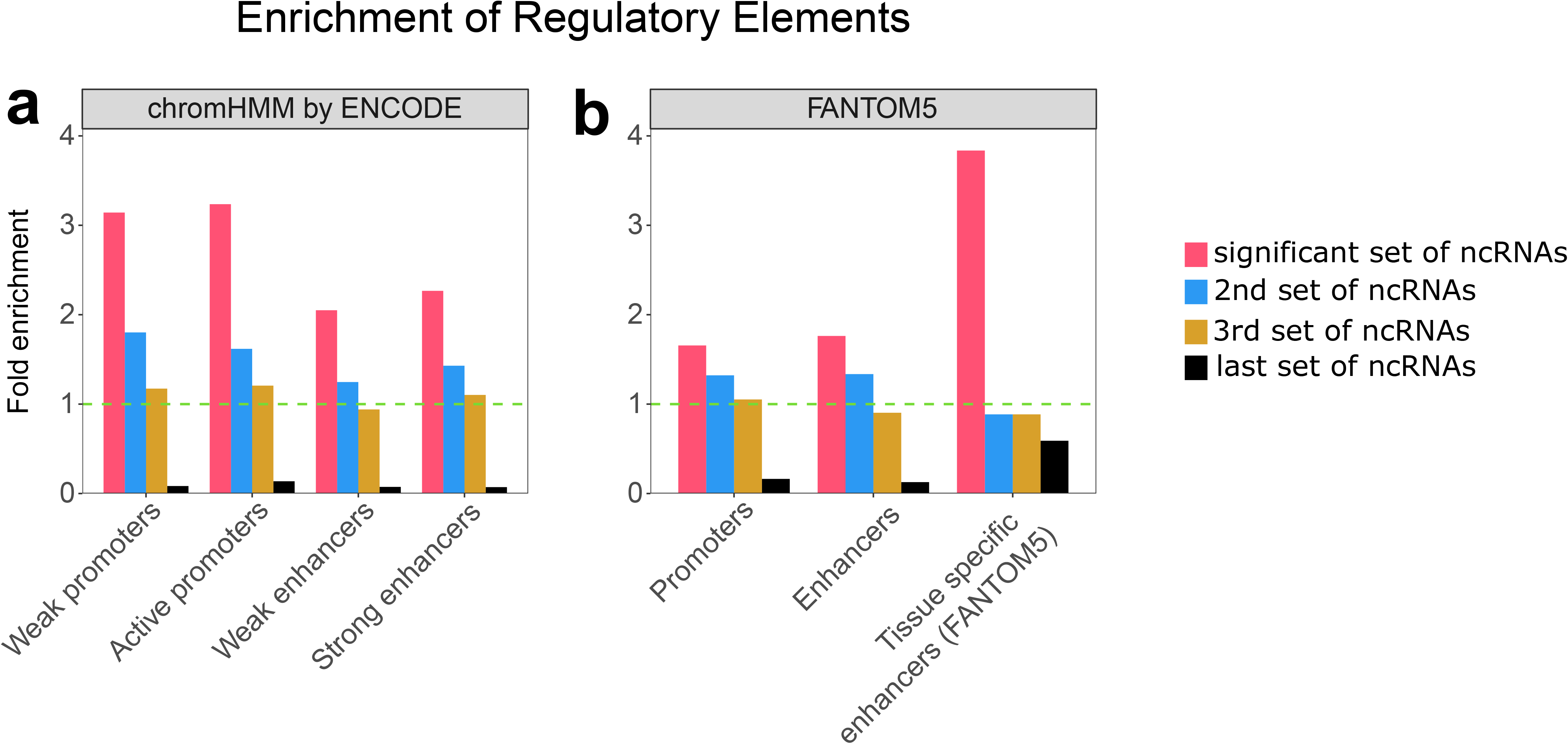
Enrichment of promoters and enhancers in the significant set of ncRNA genes. **a**) Enrichment of HMEC-related promoters and enhancers identified by ENCODE (using chromatin segmentation by HMM). **b**) Enrichment of promoters, enhancers, and breast tissue differentially expressed enhancers identified by FANTOM5. The enrichment is calculated by dividing the proportion of significantly mutated ncRNAs that overlap with each item by the proportion of all ncRNAs that overlap with that item. This enrichment is calculated for a significant set of ncRNAs (929 ncRNAs) shown in red color, 2^nd^ set (blue), 3rd set (brown) of highly mutated ncRNAs. The enrichment was also calculated for the last set of ncRNAs that had no mutation in breast cancer samples. Each set of ncRNAs contains 929 elements.

We also investigated these enrichments in the 2^nd^ and 3^rd^ sets of ncRNAs (each set contains 929 ncRNAs) with the best mutational P-values in breast cancer and the last set of ncRNAs with worse mutational P-values (*see method section*). As **Figure 3a** shows, there is the same trend of enrichment (but at much lower level) for ChromHMM predicted promoters and enhancers in the 2^nd^ and 3^rd^ sets of most mutated ncRNAs in breast cancer (**Figure 3a**). Interestingly, there is no such trend for the last set of ncRNAs (those ncRNAs with no *de novo* mutation in breast cancer samples), supporting our hypothesis that *de novo* somatic mutations are enriched in enhancer-like ncRNAs. We provided an annotated list of candidate ncRNAs with ChromHMM in **Supplementary table S2**.

The FANTOM5 consortium has released lists of transcribed human promoters and enhancers and tissue-specific transcribed enhancers of humans using CAGE (Cap Analysis of Gene Expression [26]) to study cell-type-specific enhancers. Therefore, we investigated the enrichment of FANTOM5 promoters and enhancers that overlap with the significant ncRNAs identified in this study. **Figure 3b** shows that both FANTOM5 promoters and enhancers are enriched in the candidate ncRNAs (1.66 and 1.76 times (P-value 3.59e-34 and 3.68e-27) enrichment for FANTOM5 promoters and enhancers, respectively). In other words, 52.4% of candidate ncRNAs overlapped with FANTOM5 promoters, and 36.4% of them overlapped with FANTOM5 enhancers; however, only 34.9% and 23.5% of all ncRNAs overlapped with FANTOM5 promoters and enhancers, respectively. Performing the same analysis on FANTOM5 mammary-specific enhancers demonstrated that the proportion of candidate ncRNAs that overlap with differentially expressed enhancers in the mammary epithelial cell is 3.84 times (P-value 4.03e-05) more than the genome-wide expectation (**Figure 3b**). There is also the same trend for 2^nd^ and 3^rd^ sets of ncRNAs with the best mutational P-values in breast cancer.

An annotated list of candidate ncRNAs with FANTOM5 annotations is provided in **Supplementary table S3**. For example, the pseudogene *NKAPP1* is differentially expressed in ABL1/ABL2 knockdown (shAA) breast cancer-associated cell lines [27] and downregulated in breast cancer [28]. It is also a biomarker associated with breast cancer prognosis [29, 30]. Our analysis demonstrated *NKAPP1* as one of the most significant ncRNAs with a P-value of 3.43e-06. This non-coding gene also overlapped with both FANTOM5 enhancer and ChromHMM predicted enhancer and promoter. *CATG00000062386* is another ncRNA gene that is significantly mutated in breast cancer samples. This FANTOMCAT specific ncRNA overlapped with FANTOM5 and ChromHMM enhancers and FANTOM5 mammary epithelial cell differentially expressed enhancers, indicating a potential enhancer role for this ncRNA in breast cancer.

### Histone modifications H3K27ac and H3K4me1, CTCF binding sites, and DNase hypersensitive sites are significantly enriched for BC-associated ncRNAs

We next investigated the enrichment of HMEC-specific chromatin histone active marks (e.g., H3K27ac and H3K4me1), CTCF binding sites, and DNase hypersensitive sites in the candidate ncRNAs identified in this study. These active chromatin marks are involved in many processes, including transcriptional regulation that regulates gene expression [31]. Our investigation of histone active marks demonstrated that both histone marks H3K27ac and H3K4me1 are significantly enriched 2.1 times (P-value 3.65e-57) and 1.6 times (1.90e-42) for H3K27ac and H3K4me1, respectively, in the candidate ncRNAs (**Figure 4a**). As these histone marks act as transcriptional activation and are found in both enhancer and promoter regions, our results suggest that our candidate list of ncRNAs is important for the transcriptional process in breast tissue. We performed the same analysis on histone marks H3K27me3, which is involved in the repression of transcription. Interestingly, we did not see significant enrichment (1.15 times with P-value 5.14e-02) for histone H3K27me3 within our BC-associated ncRNAs (**Figure 4a**).

**Figure 4.**
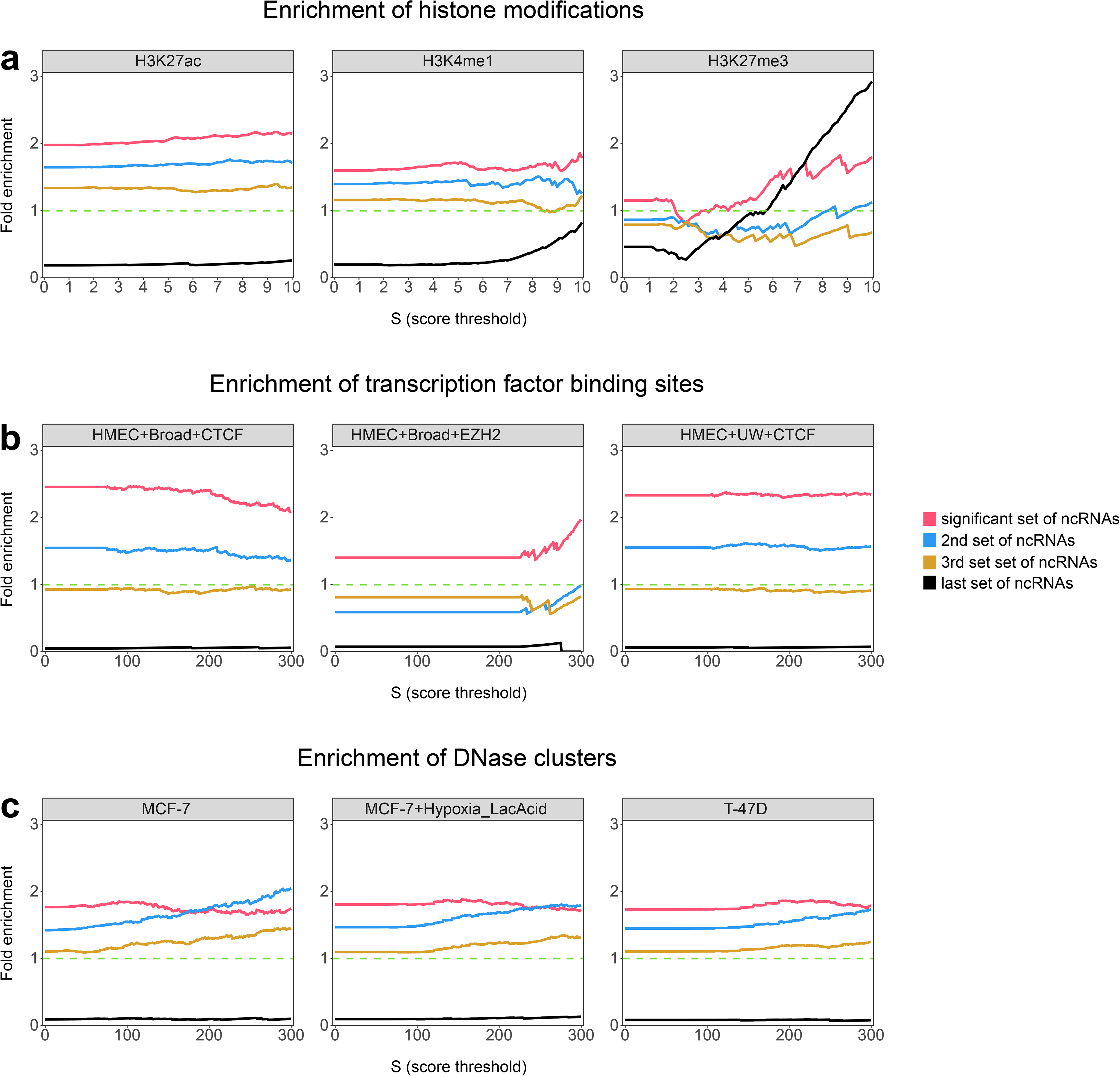
Enrichment of promoters and enhancers in the significant set of ncRNA genes. **a)** Enrichment of histone modifications (for three antibodies: H3K27ac, H3K4me1, and H3K27me3). The candidate set of ncRNAs shows significant enrichment for H3K27ac and H3K4me1 and no significant enrichment for H3K27me3, which is a repressor histone mark. **b)** Enrichment of transcription factor binding sites (for 3 ChIP-seq experiments: HMEC+Broad+CTCF, HMEC+Broad+EZH2 and HMEC+UW+CTCF). In all experiments, the ncRNAs candidate set shows significant enrichment, much higher than other sets of ncRNAs. **c)** Enrichment of DNase clusters (for three cell types: MCF-7, MCF-7+Hypoxia_LacAcid, and T-47D). As the figure shows, the candidate set of ncRNAs significantly enriched for DNase clusters, for most of the scores, much higher than other sets. As described before, the enrichment is calculated by dividing the proportion of significantly mutated ncRNAs that overlap with each item by the proportion of all ncRNAs that overlap with the item. This enrichment is calculated for a significant set of ncRNAs (929 ncRNAs) as shown in red color, 2^nd^ set (blue), 3rd set (brown) of highly mutated ncRNAs. The enrichment was also calculated for the last set of ncRNAs that had no mutation in breast cancers. Each set of ncRNAs contains 929 elements.

Transcription factors CTCF function as a transcriptional activator, repressor, insulator, or pausing transcription. In addition to CTCF sites, DNase hypersensitive sites (DHS) also have key roles in gene regulation as regulatory element markers [32]. Both CTCF and DNase are functionally related to transcriptional activity and are necessary to regulate chromatin structure. Here, we choose three CTCF ChIP-seq experiments (HMEC+Broad+CTCF, HMEC+Broad+EZH2 and HMEC+UW+CTCF) and three DHS ChIP-seq experiments (MCF-7, MCF-7+Hypoxia_LacAcid, T-47D), all related to the breast cancer. We calculated the enrichment for the four sets of ncRNAs, including significant, second, and third sets of ncRNAs with the best mutational P-values and the last set of ncRNAs with the lowest mutational P-values. Our assessment of CTCF binding and DNase hypersensitive sites also demonstrated that both CTCF (2.45 times on average with P-value 7.42e-35) and DNase (1.8 times on average with P-value 7.23e-50) are significantly enriched for our candidate ncRNA genes (**Figure 4b**). Having such significant enrichment for CTCF binding sites and DNase accessible sites is strong evidence that our significant set is not chosen randomly and is related to the gene regulation process. There is also the same trend for the 2^nd^ and 3^rd^ sets of ncRNAs (**Figure 4b**). However, the first set’s enrichments are much higher than the 2^nd^ and 3^rd^ sets of most frequently mutated ncRNAs. As **Figure 4b** shows, there is no enrichment for the last set of ncRNAs, suggesting that these ncRNAs may not be involved in the transcriptional regulation.

For example, *LINC00535* is an antisense non-coding gene that is known to be associated with breast cancer [33]. *LINC00894* is another non-coding gene that is the most downregulated lncRNA in MCF-7/TamR cells [34]. These ncRNAs are significantly mutated in breast cancer samples with P-values 4.21e-3 and 1.53e-11, respectively. *LINC00152* is another example of ncRNAs with a substantial role in enhancing breast cancer, which causes inactivation of the BRCA1/PTEN by DNA methyltransferases as tumorigenesis, mainly in triple-negative breast cancer (TNBC) [35]. All these ncRNAs overlapped with ENCODE predicted enhancers, histone 27 acetylation, CTCF binding sites, and DNase hypersensitive sites, suggesting a potential transcriptional regulatory role for these ncRNAs. An annotated list of candidate ncRNAs with these features is provided in **Supplementary tables S4 and S5**.

### BC-associated GWAS SNPs are significantly enriched in the candidate non-coding RNAs

Multiple genome-wide association studies identified disease-associated genes and their respective pathways, which provided a comprehensive understanding of the disease’s etiology. It has been reported that more than 93% of disease-associated variations found by GWAS are located in the non-coding regulatory regions of genomes [36], suggesting non-coding regulatory regions are relevant to disease and genetic mutations in gene regulatory regions is a significant mechanical contributor to diseases. To examine the enrichment of BC-associated GWAS SNPs in the candidate ncRNAs, we extracted the BC-associated SNPs from a pooled list of two GWAS datasets from the EBI GWAS Catalog [37] and GWASdb v2 from the Wang Lab [38] (*for more details see the method section*). As **Figure 5a** shows, BC-associated GWAS SNPs are significantly enriched (P-value 2.3e-27) in the candidate ncRNAs. This enrichment is much higher for those ncRNAs that contain more than 4 GWAS SNPs (> 10 times enrichment). Interestingly, when we performed the enrichment analysis for ncRNAs with more than 5 GWAS SNPs, only the candidate list of ncRNAs shows enrichment for GWAS SNPs (> 20 times), and there is no enrichment in the 2^nd^ and 3^rd^ sets of ncRNAs (**Figure 5a**). Performing the same analysis on lung cancer-associated GWAS SNPs did not show such enrichment for our candidate ncRNAs (**Figure 5a**), indicating the candidate ncRNAs are relevant to breast cancer. For example, candidate ncRNAs *RP11-353N4.6* are known to carry breast cancer associated GWAS SNPs [39]. More details on the annotated list of ncRNAs with BC-related GWAS SNPs can be found in **Supplementary table S6**.

**Figure 5.**
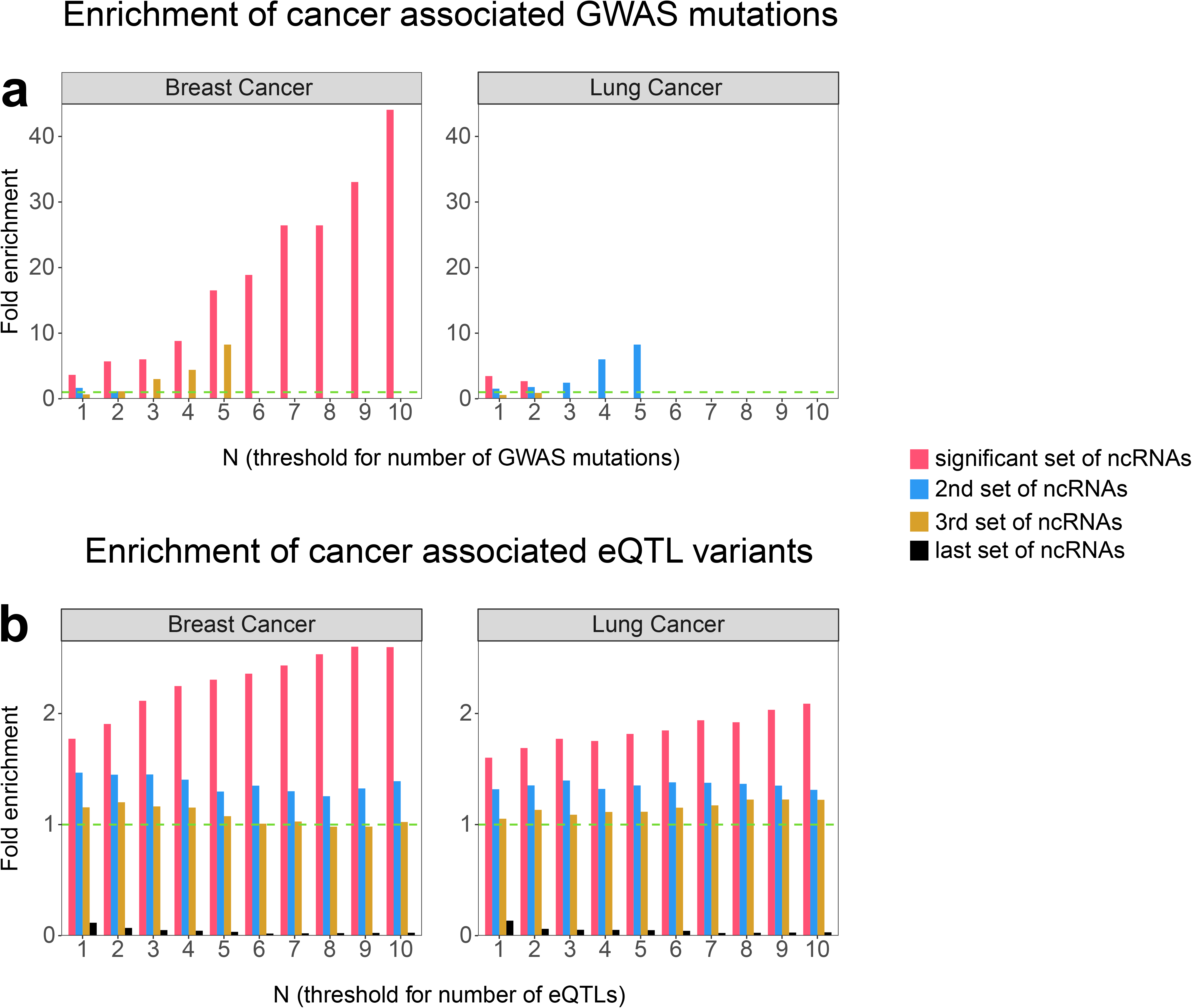
Enrichment of promoters and enhancers in the significant set of nncRNA genes. **a)** Enrichment of breast and lung cancer-associated GWAS SNPs in the candidate non-coding RNAs. **b)** Enrichment of breast and lung tissues associated with eQTL pairs in the candidate non-coding RNAs. We repeated the enrichment analysis with many items (e.g., GWAS SNP or eQTL polymorphism) overlapping the ncRNA. E.g., counting the number of ncRNAs that encompass at least 2/3/4/5/6/7/8/9/ GWAS SNPs or eQTL polymorphisms. As the figure shows, the candidate set of ncRNAs significantly enriched for both BC-related GWAS SNPs and breast tissue related eQTL polymorphisms. This enrichment is much higher than the enrichment in lung cancer related GWAS SNPs and lung tissue related eQTL polymorphisms.

### BC-associated non-coding RNAs have a significantly higher fraction of eQTL polymorphisms

Expression quantitative trait loci are genomic locations that influence gene expression in disease-related tissue and are important for understanding its mechanisms. To assess the level of eQTL polymorphisms in the candidate non-coding genes, we calculated the enrichment of breast mammary tissue eQTL polymorphisms downloaded from GTEx consortia [40] in the candidate ncRNAs. Our analysis revealed that breast mammary tissue eQTL polymorphisms are significantly enriched (1.77 times with P-value 4.11e-20) in the candidate ncRNAs. Interestingly, this enrichment is much higher for those ncRNAs that contain more than three eQTL polymorphisms (> 2x enrichment), and such an increase has not been seen in the 2^nd^ and 3^rd^ sets of ncRNAs (**Figure 5b** and **Supplementary table S6**). As a control, we calculated the enrichment of lung-specific eQTL polymorphisms in the candidate ncRNAs. As **Figure 5b** shows, the enrichment of lung-specific eQTL polymorphisms in the breast-associated ncRNAs is much lower than the enrichment observed for mammary tissue eQTL polymorphisms (**Figure 5b**). For example, lncRNA *RP11-37B2.1* and *RP11-426C22.5* are two of our candidate ncRNAs with significant P-values 7.91e-4 and 6.72e-06, respectively. These ncRNAs encompass 232 and 111 eQTL polymorphisms, respectively. lncRNA *RP11‐37B2.1* influences the risk of tuberculosis and the possible correlation with adverse drug reactions (ADRs) from tuberculosis treatment [41], and *RP11-426C22.5* is downregulated in SW1990/GZ Cells [42]. Both *RP11-37B2.1* and *RP11-426C22.5* overlapped with histone active mark H3K27ac, ChromHMM potent enhancer, and DNase hypersensitive sites, suggesting a potential transcription regulation role of these ncRNAs.

### Chromosome conformation capture data shows a potential regulatory role for BC-associated non-coding RNAs

High-throughput chromosome conformation capture (Hi-C) based assays have been used to successfully identify regulatory regions and targets of disease-associated variations [43, 44]. To gain a further understanding of the genes that our candidate ncRNAs are interacting with, we analyzed two publicly available Hi-C datasets from HMEC obtained from the Rao *et al*. study [45]. Here, we used MHiC [46] and MaxHiC [47] to analyze Hi-C raw data and identify statistically significant interactions, respectively. We identified 188,982 statistically significant interactions (P-value < 0.01 and read-count ≥ 10 – *see method section*) in the Hi-C library 1. For 6,187 of the interactions (%3.3), one side of the interaction overlapped with at least one candidate ncRNA and another side of the interaction overlapped with protein-coding genes (promoter regions of coding genes – *see method section;* **Supplementary table S7**). Repeating this analysis on the second Hi-C library also identified 318,034 statistically significant interactions. For 9,879 of the interactions (3.1%), one end of the interactions overlapped with the candidate ncRNAs and another end overlapped with the promoter region of protein-coding genes (**Supplementary table S7**). We identified 1,167 common significant interactions between the two libraries where one side of the interaction overlaps with at least one candidate ncRNA (**Supplementary table S8**) and another side with protein-coding genes. In other words, for 226 ncRNAs out of a pool of 929 candidate ncRNAs (24%), there was at least one significant interaction in both the libraries (**Supplementary table S8**); this is significantly higher (1.74 times; P-value 4.61e-11) than all ncRNAs that overlapped with one side of the interactions in both libraries (14%).

Interestingly, for 757 significant ncRNAs (%82), we identified at least one interaction in either Hi-C library 1 or library 2, resulting in 21,564 interactions. For 19,674 out of 21,564 interactions (**Supplementary table S9**), one end of the interaction that encompasses candidate ncRNAs overlapped with either ENCODE HMM predicted enhancer or active histone mark H3K27ac (both presented in HMEC). This observation suggests a potential enhancer role for these ncRNAs; In many cases, another end of the interactions overlap with protein-coding genes, including cancer-associated genes (**Supplementary table S9**). We provided a prioritized list of candidate ncRNAs that interacted with cancer-associated protein-coding genes in both the Hi-C libraries and overlapped with ENCODE HMM predicted enhancers and active histone mark H3K27ac (**Table 1**). For example, *MYC is* a BC-associated protein-coding gene, acting as a transcription factor. In common with three other transcription factors (*POU5F1*, *SOX2*, and *KLF4*), it can induce epigenetic reprogramming of somatic cells to an embryonic pluripotent state [48]. We found significant Hi-C interactions between *MYC* and two of our candidate ncRNAs *CASC8* and *PVT1* in both Hi-C libraries. *PVT1* is a known enhancer for *MYC* [49], however, *CASC8* has not yet been identified as a putative enhancer for *MYC*. There are ~200kb genomic distances between the *CASC8* and transcription start site (TSS) of MYC in which there is no protein-coding gene that overlaps with *CASC8*. There are strong signals of histone active marks and ENCODE predicted enhancers, and most importantly, FANTOM5 breast differentially expressed enhancer overlapped with *CASC8,* suggesting a potential enhancer region in *CASC8* (**Figure 6**). Interestingly, there is no somatic mutation or BC-associated GWAS SNPs overlap with *MYC*, however, *CASC8* is significantly mutated in breast cancer samples, and most importantly, it encompasses ten BC-associated GWAS SNPs. Altogether, this evidence may indicate a putative enhancer role of *CASC8* for *MYC*.

**Table 1.**
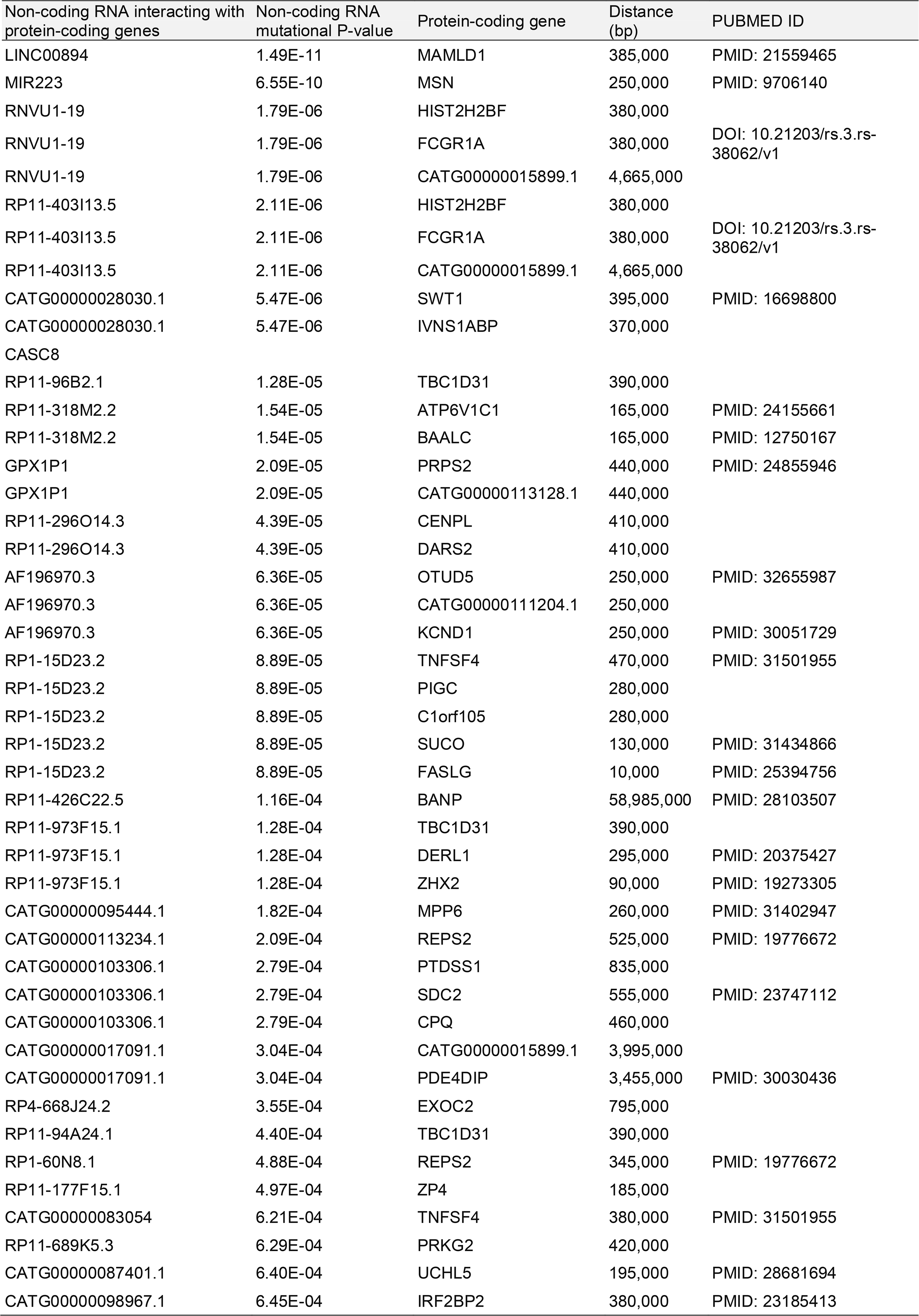
Prioritized list of candidate ncRNAs. Prioritized list of candidate ncRNAs interacting with both the Hi-C libraries and overlapped with ENCODE HMM predicted enhancer or active histone mark H3K27ac. A PUBMED ID is provided if protein-coding genes are known to be associated to cancer. A detailed list of Hi-C analyses is given in **Supplementary tables S8-S9**.

**Figure 6.**
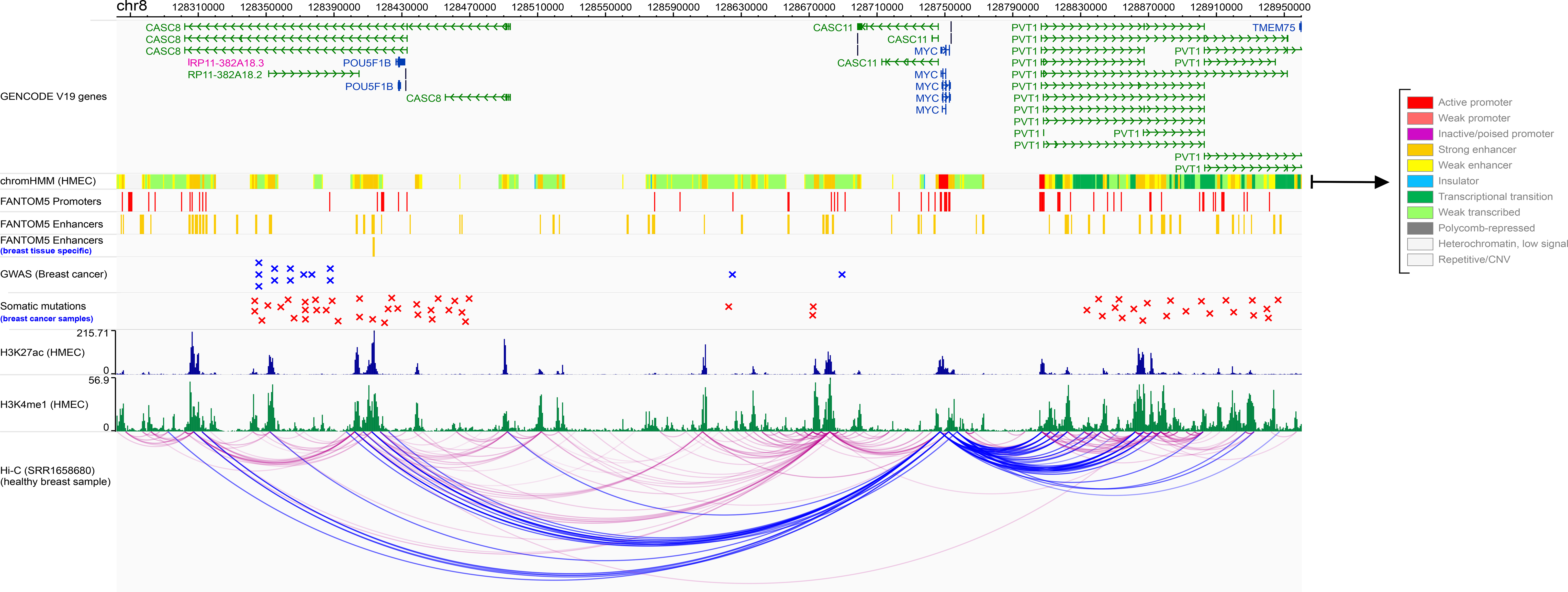
An example of Hi-C interaction between lncRNA *CASC8* and protein-coding gene *MYC*. Long non-coding RNA *CASC8* is a breast-expressed lncRNA that is significantly mutated in ICGC breast cancer samples. Our analysis on HMEC-related Hi-C data shows that this lncRNA is significantly interacting with the promoter of multiple coding genes, including *MYC*, a known breast cancer-associated gene. There are numerous strong signals of ENCODE predicted ChromHMM potent enhancers, histone active marks H3K27ac, and H3K4me1 (all presented in HMEC) that overlap with *CASC8*. Importantly, FANTOM5 breast differentially expressed enhancers also overlapped with *CASC8*. Our analysis of GWAS SNPs and *de Novo* somatic point mutations revealed that *CASC8* covered multiple breast cancer GWAS SNPs and many somatic point mutations related to breast cancer samples. In contrast, *MYC* does not cover either GWAS SNP or BC-related somatic mutations. The figure also shows that the *MYC* gene interacts with *PVT1,* another significantly mutated lncRNA in breast cancer sample (30kb far away from *MYC*). *PVT1* is also overlapped with breast tissue-related regulatory features and is a previously known enhancer for the *MYC* gene. We used MHiC [46] to analyze raw Hi-C data and MaxHiC [47] to identify significant Hi-C interactions. Significant interactions are shown in blue color. ENCODE predicted chromHMM files chromatin signals were used in peak format.

Another example is non-coding RNA *CATG00000061359* is a FANTOMCAT-specific intergenic lncRNA that is significantly mutated in breast cancer samples (P-value 5.32e-4). There is a significant Hi-C interaction between *CATG00000061359* and gene *GTPBP8* (previously associated with breast cancer [50]) in both the libraries. We also found a breast tissue-associated eQTL polymorphism in this lncRNA that influences the expression of *GTPBP8* in breast tissue. Interestingly, both HMEC specific H3K27ac and HMM predicted enhancer overlap with this lncRNA. This may suggest that variation in *CATG00000061359* may influence the expression of *GTPBP8* in breast cancer. We have provided an annotated list of candidate ncRNAs with Hi-C interactions in **Supplementary table S9**.

## Discussion

Somatic point mutations play a key role in tumorigenesis and development of cancer [51]. Recent studies on somatic mutation evolution in cancer have led to the identification of cancer driver genes [52] and cancer mutational signatures [53, 54]; However, the analysis of somatic mutations has focused mainly on the protein-coding genes of the genome and their potential contribution in the ncRNAs have been far less studied, particularly their functional significance across long ncRNAs. Recent studies have uncovered that the ncRNAs are involved in breast cancer development through a variety of mechanisms including regulating the expression of protein-coding genes and functions at transcriptional, translational, and post-translational levels [18–22]. In this study, we first focused on identifying ncRNAs that were significantly and specifically mutated in breast cancer patients and then uncovered the connection between somatic point mutations in BC-associated ncRNAs and ncRNA regulatory properties in breast cancer. We have shown that the candidate lncRNAs with enrichment of somatic point mutations have a much higher fraction of regulatory features compared to genome-wide expectation, suggesting the potential impact of somatic mutations on the regulatory function of lncRNAs. Notably, we have shown that most of the candidate lncRNAs are interacting with promoters of protein-coding genes, again indicating the potential regulatory role of lncRNAs with significant enrichment of somatic point mutations in breast cancer. Our analysis demonstrated that highly mutated lncRNAs have more cell type-specific functions and are more likely to interact with protein-coding genes. To our knowledge, this association has not been previously explored. However, our analysis is limited to the associations identified by pure data driven analyses. Future experimental validation can reveal finer details about the interdependence of lncRNA function in breast cancer.

We have developed an extensive web-based resource to communicate our results with research community. Further works could focus on the candidate lncRNAs provided in this resource to check their complex regulatory functions and reveal novel mechanism underlying carcinogenesis and breast cancer treatment.

We also presented a novel computational method, SomaGene, to prioritize and predict disease associated ncRNAs by integrating multi-level omics data including somatic mutations, transcriptomic and epigenetic signals, and chromatin confirmation data.

## Conclusion

NcRNAs have long been considered as non-functional part of the human genome [55], however, these non-coding elements (majority lncRNAs) have recently opened a new insight in the study of breast cancer, acting as indispensable contributors to many cellular activities including the proliferation, apoptosis, survival, differentiation and the breast cancer metastasis [56]. In addition, lncRNAs have been used as biomarkers in many cancers, including breast cancer, indicating their potential in diagnosis, prognosis, and therapeutics. In this study, we performed an integrative analysis to present the first comprehensive resource (http://www.ncrna.ictic.sharif.edu and https://www.ihealthe.unsw.edu.au/research) of ncRNAs that significantly and specifically mutated in breast cancer genomes. We showed that the majority of those ncRNAs act as enhancers, suggesting enhancer mutations may be relevant to the functions for such ncRNAs in breast cancer. Our results provide an extensive resource for researchers that can be used in the future to better understand the underlying genetic risk factors that contribute to breast cancer. We believe our study sets the stage for a new framework for future research in the role of non-coding RNAs in the treatment of breast cancer. We also developed SomaGene as a user-friendly tool to be used in the analysis of non-coding RNA genes in all cancers.

## Methods and Materials

### Tool availability

The SomaGene open-source R package, a sample dataset, and instructions on how to run SomaGene are provided at https://github.com/bcb-sut/SomaGene.

### ICGC Dataset

We used the ICGC dataset, which contains somatic point mutations from 1,855 breast cancer samples and 10,419 samples from 19 different types of cancers.

### A combined list of non-coding RNAs from FANTOMCAT and Ensembl consortia

We have combined two genes lists from FANTOM5 [26] and Ensembl [57] consortia (genome building hg19) that enables us to have a comprehensive list of non-coding RNA genes. We used an in-house script to combine the lists based on gene coordinates and/or gene names. If both consortiums have the same gene but different gene coordinates, we considered the FANTOM5 genes as the priority. In total, 60,156 non-coding RNA genes were available in the combined list.

### Identify significantly mutated non-coding genes

We included all samples from ICGC with at least one *de novo* mutation. We only considered single nucleotide mutations and excluded insertions or deletions from the analyses. To identify ncRNAs that significantly mutated in breast cancer samples compared to other cancers, we used Fisher’s exact test and permutation testing in the following manner:

We calculated a P-value for each ncRNA using one-sided Fisher’s exact test applied to a 2×2 contingency table whose elements are (1) the number of samples in breast cancer that are mutated in this ncRNA, (2) the number of samples in breast cancer that are not mutated in this ncRNA, (3) the number of samples in all cancers other than breast cancer that are mutated in this ncRNA and (4) the number of samples in all cancers other than breast cancer that are not mutated in this ncRNA (**Supplementary figure S2**). To identify significant ncRNAs, we calculated P-values for 1,000,000 random permutations (define by user) of sample IDs across all cancers to estimate the probability that an association emerges by chance confidence interval of 99%. From this, we identified a total of 929 significantly mutated ncRNAs in breast cancer.

### Overlap and aggregation score methods

To adequately examine the overlapping of ncRNAs with regulatory features and calculate the overlapping score with each element, each annotation’s overlapping ranges were used in our analysis. In the case of FANTOM5 promoters, enhancers, and tissue-specific enhancers, three binary variables for each ncRNA were calculated, indicating that an ncRNA has overlapped with any enhancer or promoter in the FANTOM5 dataset (**Supplementary table S2**). In eQTL annotation, a set of all entries whose location overlaps with each ncRNA was extracted by concatenating the variation_id and gene_id for each location.

The chromatin segmentation annotation comprises genomic ranges, which each of them attributed to one of 11 functional categories (segments). The percentage of overlap with a segment was calculated as the sum of proportions of nucleotides in that segment’s ranges that overlapped with the ncRNA. Thus, a profile of overlapping chromatin segments was calculated with their corresponding coverage over the ncRNA (**Supplementary figure S3, Supplementary table S4**).

For histone modifications annotation, besides the proportions of nucleotides covered with the overlapping histone modification ranges in each ncRNA, the peak scores of these ranges were averaged together by the corresponding overlap percentages as their weights to obtain a single histone modification score. The coverage (overlap percentage) of each ncRNA with histone modification ranges was also calculated as the total proportion of nucleotides in the ncRNA covered with histone modification ranges.

In the case of DNase annotation, each genomic range is attributed to a group of cell types that show DNase hypersensitivity in that area. For each range, several cell type IDs with their corresponding DNase hypersensitivity scores are associated. We combined the annotation ranges that overlap with it to get an aggregated set of cell-type IDs, scores, and overlap percentages by calculating the proportion of nucleotides in the ncRNA covered with these ranges. After this step, several IDs were duplicated in many aggregated sets that were summed over the overlap percentages to obtain a single overlap measure. Also, it was averaged over its scores by the corresponding overlap percentages as their weights to take a single score for that ID. For each genomic range in transcription factors annotation, a group of transcription factors (from ENCODE) with their corresponding ChIP-Seq peaks are reported. While the schema of transcription factors annotation is similar to that of DNase annotation, the same procedure as described for DNase annotation was performed to obtain an aggregated set of transcription factor IDs, scores, and overlapping percentages for each ncRNA (**Supplementary table S5**). The overlap of BC-related GWAS mutations loci with the overlapped ncRNA regions was identified. The total number of overlaps for each ncRNA was recorded in the output table (**Supplementary table S6**).

### Calculating enrichments

Every “enrichment” that is calculated throughout this study is defined as “the fraction of ncRNAs in the significant set that have the trait of interest (e.g., having overlap with or having a minimum score of a specific annotation) divided by genome-wide expectation”. It can be easily justified that the mentioned value equals the fraction of items in the whole set of ncRNAs with the trait of interest: assume that the total number of ncRNAs is ***N***. The number of significant ncRNAs is ***S*** while ***A*** items among all and ***Y*** items within the significant set have the property of interest. If a random collection of ncRNAs with size ***S*** is sampled, then the probability that ***X*** items within this set happen to have the property of interest follows a binomial distribution with parameters ***S*** and ***A/N***, i.e., ***X*** ~ *Binom(**S, A/N**)*. Thus, the expected value of ***X*** equals ***S*** × ***A/N*** which shows that the expected fraction of items having the property of interest in a random set of ncRNAs with size ***S*** equals ***A/N***. We then conclude that the defined enrichment can be calculated as (***Y/S***)/(***A/N***).

### FANTOM5 promoters, enhancers, and breast differentially expressed enhancers

The online FANTOM5 CAGE expression atlas [24] represents the transcription of many cell-types states such as promoter, enhancers, and gene regulation in primary human cells, tissues, and cell lines. We used FANTOM5 CAGE expression atlas to interrogate significant ncRNAs to identify a set of significant ncRNA that overlapped with promoter and enhancer independently. The entire collection of enhancers and promoters found in the FANTOM5 data were downloaded from the FANTOM5 Phase2 [26, 58]. We also downloaded FANTOM5 breast differentially expressed enhancers from [59].

### ENCODE chromatin state segmentation

The Chromatin State Segmentation uses a standard set of states that learned by computationally integrating ChIP-Seq data for nine factors plus input using a Hidden Markov Model (HMM) across various cell types. Also, it shows a classification of chromatin, like “enhancer,” “promoter,” or “repressed.” We used this dataset to interrogate with ncRNAs to identify the set of breast-specific candidate ncRNAs overlapped with chromatin state. The complete set of chromatin state segmentation for the models derived from HMECs, grown in vitro related to breast cancer, was downloaded from the ENCODE project [60].

### ENCODE histone modifications by ChIP-Seq

For this study, we used HMEC-specific ChIP-Seq data in the form of processed peak calls for histone modifications H3K27ac, H3K27me1, H3K4me3, and CTCF from the ENCODE project to interrogate with significant ncRNAs to identify a set of significant ncRNA that overlapped with three types of histone modifications [60].

### ENCODE transcription factor ChIP-Seq data

This data shows regions of transcription factor binding derived from an extensive collection of ChIP-Seq experiments and DNA binding motifs identified within these regions by the ENCODE Factorbook repository [61]. The transcription factors and proteins are responsible for modulating gene transcription bind as assayed by chromatin immunoprecipitation with antibodies specific to the transcription factor followed by sequencing of the precipitated DNA (ChIP-Seq). We used this dataset to interrogate significant ncRNAs to identify a set of significant ncRNA that overlapped with transcription factor ChIP-Seq. The set of transcription factor ChIP-Seq data for the HMEC cell line were downloaded from ENCODE project [60].

### DNase clusters

Regulatory regions tend to be DNase-sensitive which are accessible chromatin zones and functionally related to transcriptional activity. We used DNase hypersensitive sites from MCF-7 and T47D from ENCODE project to interrogate significant ncRNAs to identify a set of significant ncRNA that overlapped with DNase clusters.

### Hi-C data analysis

Hi-C is an experiment for identifying the number of interactions between genomic loci near 3D space. In our study, we used two replications related to the HMEC [45]. We used MHiC [46] and Hi-C Pro [62] with the default parameters for analyzing and aligning Hi-C data in 5kb fragment size. We then used MaxHiC [47] as a background correction model to identify significant Hi-C interactions for getting true cis-interaction. Here, we included those significant interactions with P-value < 0.01, read-count >= 10, with the distance between two sides of interaction more than 5kb and less than 20Mb. We then annotated Hi-C interactions with coding and non-coding genes from our combined genes list. At least 10% overlap between gene and Hi-C fragments has been considered to annotate Hi-C fragments with genes.

### Genotype-tissue expression eQTLs

eQTL data were downloaded from the Genotype-Tissue Expression (GTEx) project [40]. We used the set of GTEx v7 eQTLs identified as significant in the HMEC line from the GTEx project.

### Literature search for non-coding interacting genes

Our literature searches were focused on human studies and English language publications available in the PubMed, Scopus, and Web of Science. Both Medical Subject Headings (MeSH) terms and related free words were used in order to increase the sensitivity of the search. We also used data and text mining techniques to extract additional related studies [63–69]. A decision tree approach and a knowledge-based filtering system technique have been also used to categorize the texts from the literatures search [67, 70]. The search terms included “noncoding RNA” or “lncRNAs” or “genes name + cancer”. “BC” or “breast carcinoma” and “breast neoplasm”.

### Genome-wide association study (GWAS) analysis

We have pooled two GWAS datasets, EBI GWAS Catalog [37] and GWASdb v2, from Wang Lab [38] to interrogate significant ncRNA to identify a set of significant ncRNAs that overlapped with GWAS SNPs. We first converted all GWAS SNP coordinates to UCSC hg19 using UCSC Lift Genome Annotations tools [71] *(www.ncbi.nlm.nih.gov/genome/tools/remap)* and then used an in-house script to combine both GWAS datasets based on gene coordinates and/or gene symbols.

## Author contributions

HAR designed the study; HAR, NR, and MB wrote the paper. HAR, NR, JB, MST, MB, NHL and HRR edited the manuscript. NR, MB, MH carried out all the analyses, including the statistical analyses, gene prioritization, annotation, and permutation, Hi-C data analysis with the supervision of HAR and HRR. NR and MB generated all figures and all tables with the supervision of HAR and HRR. SK designed and developed the website. All authors have read and approved the final version of the paper.

## Data and tool availability

The source code and a sample dataset can be accessed at https://github.com/bcb-sut/SomaGene.

## Acknowledgments

We kindly acknowledge the Government of Western Australia, Department of Health, Clinical Excellence, for their kind support on this project through the MERIT award to H.A.R. H.A.R is also supported by UNSW Scientia Program Fellowship. Analysis was made possible with computational resources provided by the BioMedical Machine Learning Bioinformatics Server with funding from the Australian Government and the UNSW SYDNEY. H.R.R is supported by IRN National Science Foundation (INSF) Grant No. 96006077.

## Supplementary figures legends

**Figure S1.**
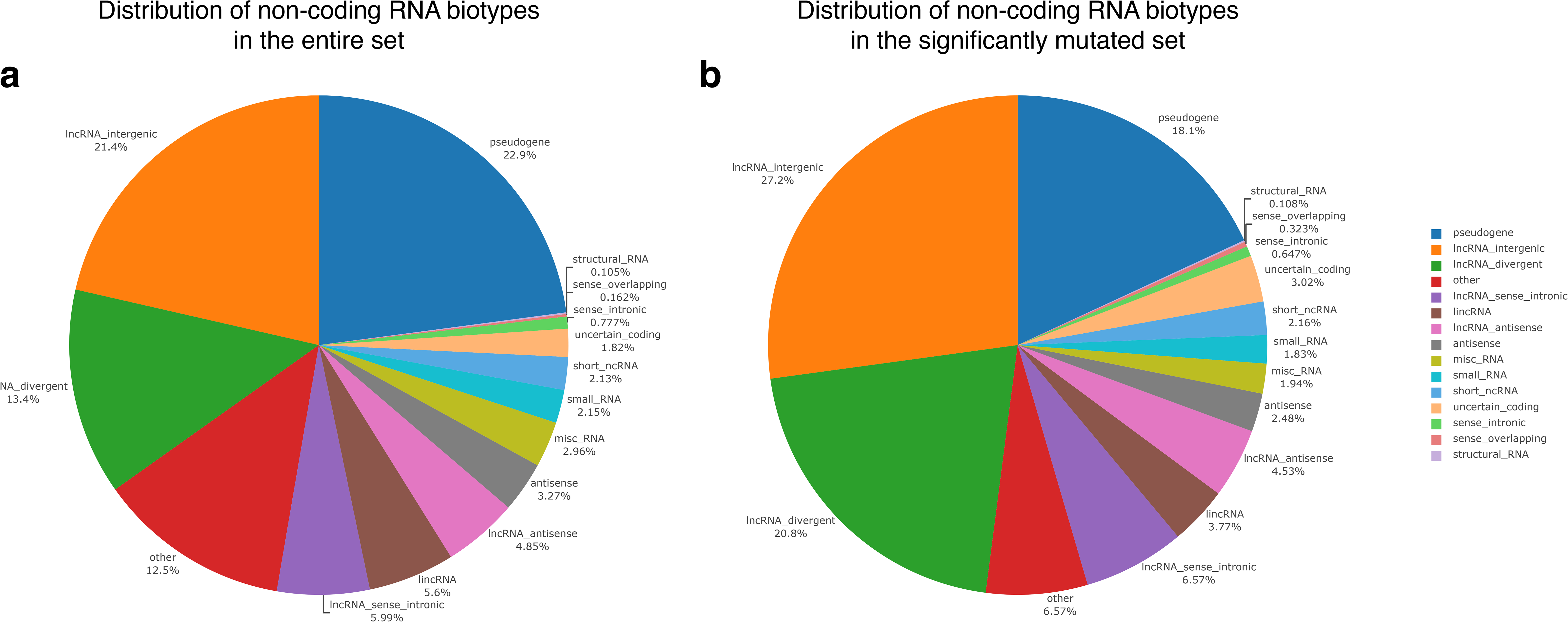
The proportion of classes of ncRNAs. **a)** for the entire dataset. **b)** for the set of significantly mutated non-coding RNAs.

**Figure S2.**
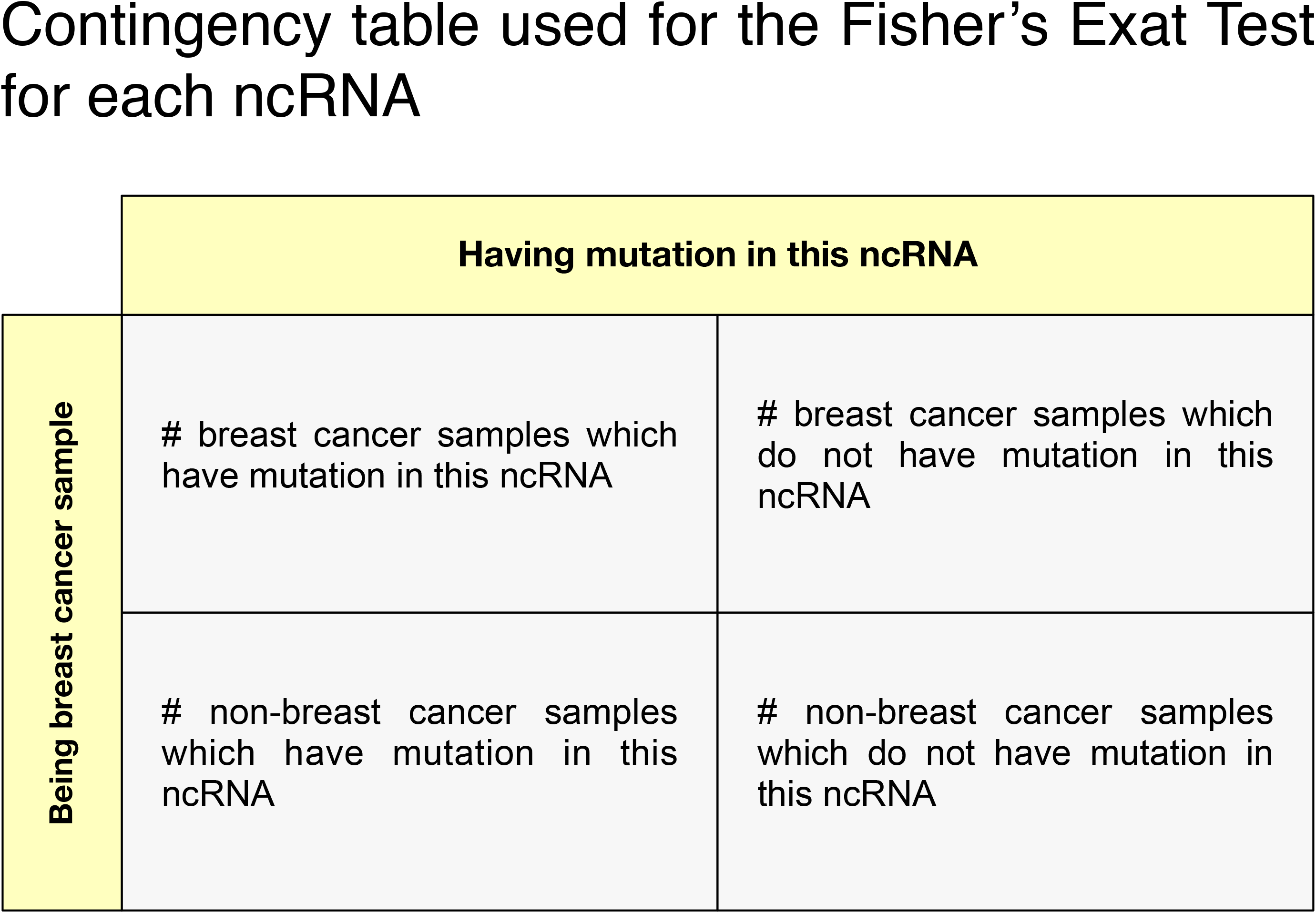
The contingency table was used for Fisher’s exact tests. For each ncRNA, a contingency table is constructed based on this scheme to perform a Fisher’s exact test to determine if the corresponding non-coding gene is significantly mutated in breast cancer samples compared to other cancer types.

**Figure S3.**
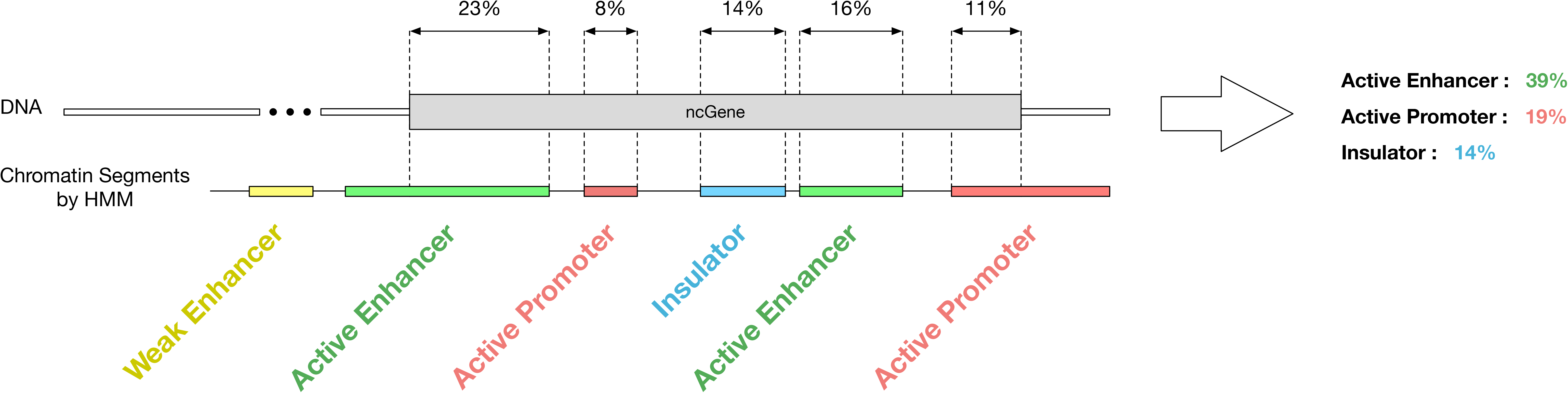
Calculation of overlap between a non-coding gene and chromatin segments. This diagram shows an example situation in which a non-coding gene overlaps with different types of segments in chromHMM annotation. The percentage of overlap with each segment is calculated as the total number of nucleotides in the non-coding gene region covered by each type of segment divided by the non-coding gene length.

## Supplementary tables legend

**Table S1. Significantly mutated ncRNAs.** A list of non-coding RNAs which are significantly mutated in breast cancer samples compared to samples with other cancers (P-value < 0.05).

**Table S2. Annotate candidate ncRNAs with ENCODE chromHMM predicted chromatin states.** Annotating significantly mutated ncRNAs (candidate ncRNAs) with ENCODE predicted chromatin state marks (chromHMM) presented in the HMEC cell line.

**Table S3. Annotate candidate ncRNAs with FANTOM5 features.** Annotating significantly mutated ncRNAs (candidate ncRNAs) with FANTOM5 promoters, enhancers, and FANTOM5 breast differentially expressed enhancers.

**Table S4. Annotate candidate ncRNAs with histone marks.** Annotating significantly mutated ncRNAs (candidate ncRNAs) with HMEC related histone modification, including CTCF, H3K27ac, H3K4me1, and H3K4me3. For each category and each gene, we identified the percentage of overlap and average score.

**Table S5. Annotate candidate ncRNAs with DNase hypersensitive sites and transcription factors.** The list of significantly mutated ncRNAs (candidate ncRNAs) with HMEC-related DNase hypersensitive sites (DHSs) and transcription factors. For each gene and each category, we calculated the percentage and score of the overlapping. The name of each category is shown in the “Details for DNase Clusters” sheet.

**Table S6. Annotate candidate ncRNAs with BC-associated GWAS SNPs and breast tissue-related eQTL polymorphisms.** Annotating significantly mutated ncRNAs with breast cancer-related GWAS SNPs. GWAS SNPs were downloaded from EBI GWAS Catalog [37] and GWASdb v2 from Wang Lab [38]. All GWAS SNPs with a P-value less than 1e-8 were considered. We also used GTEx breast-related eQTL polymorphisms to identify how many of our candidate ncRNAs encompass at least one breast tissue-related eQTL polymorphism. We calculated the number of eQTL polymorphisms and the eQTL paired coding genes for each ncRNA.

**Table S7. Annotate ncRNAs with HMEC related Hi-C interactions.** We used chromosome conformation capture data (Hi-C) as evidence of potential enhancer activity for our candidate ncRNAs. We used two replications of HMEC-related Hi-C data from the Rao *et al.* study [45]. We only reported significantly interacting regions for each replication. We identified which ncRNA and coding gene overlapped with interaction for each fragment and annotated the ncRNA side of the interactions with breast-related regulatory features.

**Table S8. Same as Table S7, but only common interactions are considered.** We only considered those Hi-C interactions that appeared as significant interactions in both replications (common interactions between SRR1658680 and SRR1658686-9). Common interactions also filtered out if neither left nor right sides of the interaction overlapped with our candidate ncRNAs.

**Table S9. Same as Table S8, but we considered those interactions that become significant in either Hi-C replicate1 or replicate 2.** We considered those Hi-C interactions that appeared as significant interactions in at least one of the replications. Significant interactions also filtered out if neither left nor right sides of the interaction overlapped with our candidate ncRNAs.

